# Microglia-specific NF-κB signaling is a critical regulator of prion-induced glial inflammation and neuronal loss

**DOI:** 10.1101/2024.09.12.612597

**Authors:** Arielle J. D. Hay, Katriana A. Popichak, Genova Mumford, Payton Shirley, Jifeng Bian, Lauren Wolfrath, Samantha Lei, Michael Eggers, Eric M. Nicholson, Ronald B. Tjalkens, Mark D. Zabel, Julie A. Moreno

## Abstract

Prion diseases are a group of rare and fatal neurodegenerative diseases caused by the cellular prion protein, PrP^C^, misfolding into the infectious form, PrP^Sc^, which forms aggregates in the brain. This leads to activation of glial cells, neuroinflammation, and irreversible neuronal loss, however, the role of glial cells in prion disease pathogenesis and neurotoxicity is poorly understood. Microglia can phagocytose PrP^Sc^, leading to the release of inflammatory signaling molecules, which subsequently induce astrocyte reactivity. Animal models show highly upregulated inflammatory molecules that are a product of the Nuclear Factor-kappa B (NF-κB) signaling pathway, suggesting that this is a key regulator of inflammation in the prion-infected brain. The activation of the IκB kinase complex (IKK) by cellular stress signals is critical for NF-κB-induced transcription of a variety of genes, including pro-inflammatory cytokines and chemokines, and regulators of protein homeostasis and cell survival. However, the contribution of microglial IKK and NF-κB signaling in the prion-infected brain has not been evaluated. Here, we characterize a primary mixed glial cell model containing wild-type (WT) astrocytes and IKK knock-out (KO) microglia. We show that, when exposed to prion-infected brain homogenates, NF-κB-associated genes are significantly downregulated in mixed glial cultures containing IKK KO microglia. Mice with IKK KO microglia show rapid disease progression when intracranially infected with prions, including an increase in microglia and reactive astrocytes, and accelerated loss of hippocampal neurons and associated behavioral deficits. These animals display clinical signs of prion disease early and have a 22% shorter life expectancy compared to infected wild-type mice. Intriguingly, PrP^Sc^ accumulation was significantly lower in the brains of infected animals with IKK KO microglia compared to age-matched controls, suggesting that accelerated disease is independent of PrP^Sc^ accumulation, highlighting a glial-specific pathology.

Conversely, primary mixed glia with IKK KO microglia have significantly more PrP^Sc^ accumulation when exposed to infected brain homogenates. Together, these findings present a critical role in NF-κB signaling from microglia in host protection suggesting that microglial IKK may be involved in sufficient clearance of prions.

## Introduction

Nuclear Factor-kappa B (NF-κB) signaling in microglia modulates inflammation and protein accumulation in a variety of neurodegenerative diseases (1-3). However, the role of this signaling pathway in microglia has not been established in prion disease. Prion diseases are rare and fatal neurodegenerative diseases that affect a variety of mammalian species, including humans. The hallmark of these diseases is the misfolding of the cellular prion protein, PrP^C^, to the transmissible form, PrP^Sc^, which acts as a seed and spreads throughout the brain, leading to neuroinflammation and irreversible neurodegeneration. Glial cells, particularly astrocytes and microglia, respond to PrP^Sc^ and release inflammatory molecules that contribute to neuronal death (4-7). The brains of prion-infected mice have increased cytokines and chemokines that are associated with the NF-κB pathway, some of which are detectable well in advance of clinical signs of disease (8-11). Previous studies have had conflicting results in whether NF-κB plays a significant role in prion pathogenesis (12, 13). These studies have knocked out critical genes in the NF-κB pathway in astrocytes and neurons, but have largely ignored the role of microglia-specific NF-κB signaling. Microglia are critical for inducing inflammatory activation in astrocytes, which in turn can produce neurotoxic signals that contribute to neurodegeneration (14-17). Moreover, when exposed to PrP^Sc^ *in vitro*, microglia respond by upregulating genes associated with NF-κB signaling (18).

The NF-κB pathway is responsible for the expression of a large number of genes, including cytokines, chemokines, enzymes, receptors, and regulators of apoptosis. Under normal conditions, NF-κB remains sequestered in the cytoplasm, inhibited from translocating to the nucleus by IκB and associated proteins. The IκB kinase complex (IKK) is activated by a variety of stimuli such as cytokines, growth factors, pathogen-associated molecular patterns (PAMPs) and damage-associated molecular patterns (DAMPs)(19, 20). In prion disease, IKK activation typically occurs through toll-like receptors (TLRs) and nucleotide-binding oligomerization-domain protein-like receptors (NLRs)(20, 21). Activated IKK phosphorylates IκBα, leading to IκBα ubiquitination and degradation by the proteosome. Uninhibited, NF-κB translocates to the nucleus and upregulates pro-inflammatory cytokines.

Here, we demonstrate that microglia-specific knock-out (KO) of IKK leads to significant changes in gene expression in primary mixed glia exposed to prion-infected brain homogenates. Prion exposed mixed glia containing both astrocytes and microglia show a drastic downregulation of NF-κB-associated genes in the absence of microglial IKK, compared to wild-type (WT) mixed glia. Intriguingly, this is not associated with a decrease in neurotoxicity. We hypothesize that this is due to the increased accumulation of PrP^Sc^ in glial cultures containing IKK KO microglia, potentially due to the role of IKK in autophagy (3, 22). We further investigated this phenomenon in mice that have myeloid cell-specific IKK KO (including macrophages, monocytes and microglia) (3, 23). These mice were inoculated intracranially with mouse-adapted scrapie prions and assessed for clinical and behavioral changes, survival, glial cell number and morphology, neuronal health and prion accumulation in the brain. Mice with IKK KO microglia succumbed to prion infection 22% faster than WT mice. Compared to age-matched WT mice, mice with IKK KO microglia showed an increase in microglia and reactive astrocytes, increased spongiosis, and a decrease in hippocampal neurons, but lower levels of PrP^Sc^ throughout the brain. Together, these findings suggest that microglial IKK and NF-κB signaling are critical for host protection against prion infection. Understanding inflammatory signaling pathways and cellular cross-talk in the prion-infected brain is critical for the development of therapeutics.

## Results

### NF-κB-associated genes are significantly downregulated in mixed glial cultures containing IKK KO microglia

To assess how IKK KO microglia respond to the milieu in the prion-infected brain, mixed glial cultures containing both astrocytes and microglia were treated with 0.1% Normal Brain Homogenate (NBH) or Rocky Mountain Laboratories (RML) mouse-adapted scrapie brain homogenates for 7 days. We previously demonstrated that RML inoculum at low concentration (0.1%) is sufficient to induce infection and detect newly synthesized PrP^Sc^, not from residual brain homogenate (24). Mixed glial cultures contain astrocytes and microglia, as demonstrated by GFAP expression, regardless of inoculum. However, Iba1+ microglia were only detectable by western blot in cultures treated with RML (Supplemental Fig 1).

RNA isolated from RML-exposed WT and IKK KO mixed glia was analyzed using a mouse NF-κB signaling pathway panel containing 84 genes of interest. A heat map analysis to of NF-κB-regulated gene expression profile demonstrates that the majority of NF-κB-associated genes are downregulated in IKK KO cultures compared to WT cultures, regardless of NBH- or RML-treatment (Fig 1A). A modest increase in NF-κB-associated genes was observed in RML-infected WT glia compared to NBH-treated WT glia (Fig 1C-F and J), consistent with previous findings (24). A substantial difference in gene expression was observed between RML-infected WT and microglia-specific IKK KO mixed glia, while IKK KO glia show a negative fold change of -1.5 or greater for 57 of the 84 genes analyzed (68%) (Fig 1A). The greatest fold changes were seen in toll-like receptor 9 (TLR9) (-193) and Bcl2a1a (-199). Few genes increased, but the largest increases were seen in epidermal growth factor receptor (Egfr) (+2.9) and macrophage colony-stimulating factor 1 (CSF1) (+2.4), involved in cell growth and microglia proliferation, respectively (25, 26).

**Fig 1.**
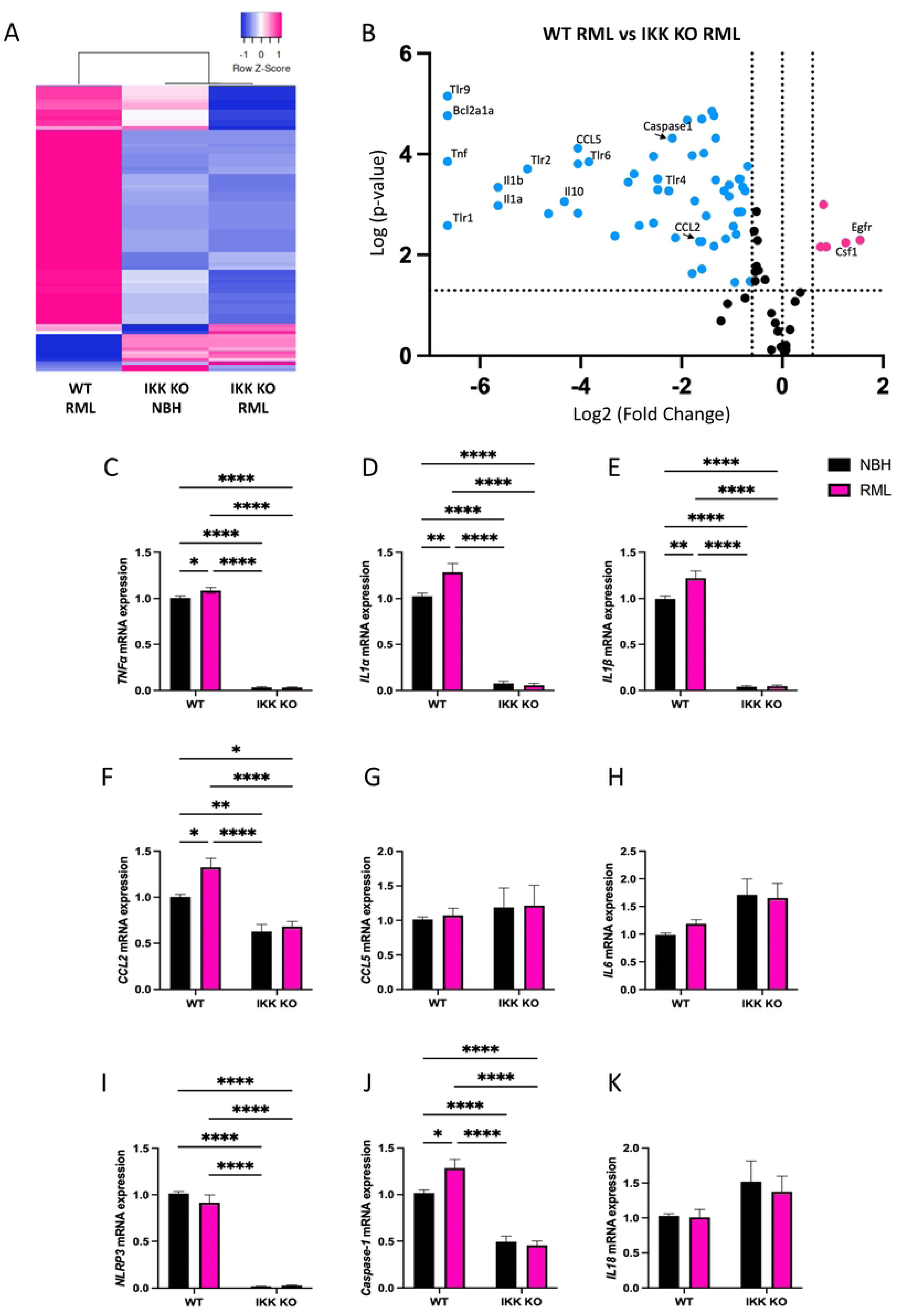
NF-κB-associated genes are significantly downregulated in infected mixed glial cultures containing IKK KO microglia. **A** Heat map comparing NF-κB-associated gene expression in infected WT mixed glia and mixed glia containing IKK KO microglia treated with NBH or RML-scrapie for 7 days, all compared to NBH-treated WT mixed glia. **B** Volcano plot comparing NF-κB-associated gene expression between RML-infected WT to microglia-specific IKK KO mixed glia shows the majority of genes are downregulated in IKK KO cultures. X-axis intersections: fold change +/- 1.5. Y axis intersection: *p*-value < 0.05. Samples are composed of 3 pooled technical replicates, and 3 biological replicates for each group. mRNA expression of NF-κB-associated cytokines and chemokines was further assessed for **C** *TNFα*, **D** *IL1α*, **E** *IL1β*, **F** *CCL2*, **G** *CCL5* and **H** *IL6* were analyzed, as were NLRP3-associated genes **I** *NLRP3*, **J** *Caspase-1* and **K** *IL18*. Analysis of 3-5 biological replicates, each with 3 technical replicates. One-way ANOVA and post-hoc Tukey test, error bars = SEM, \**p* < 0.05, \*\**p* < 0.01, \*\*\*\**p* < 0.0001.

To best observe changes in gene expression between RML-infected WT and IKK KO cultures, analysis presented in a volcano plot highlights specific genes which show significance in pathways involved in prion disease (Fig 1B). Of note, many inflammatory cytokines and chemokines are downregulated in IKK KO cultures (*tnfα*, *il1α*, *il1β*, *ccl2* and *ccl5*), as well as the anti-inflammatory cytokine *il10*. Interestingly, the NLRP3-associated gene *caspase-1* is significantly decreased. Of note, many TLRs, including *TLR1*, *TLR2*, *TLR4*, *TLR6* and *TLR9*, are all decreased in IKK KO cultures. A full list of genes with a significant fold change of + or – 1.5 is available in Supplemental Table 1.

We selected to analyze a few specific NF-κB-associated inflammatory genes that are known to be upregulated in the prion-infected brain (8, 9, 27). Additionally, we wanted to look at downstream effects on NLRP3-associated genes. Mixed glia containing WT or microglia-specific IKK KO were treated for 7 days with NBH or RML-scrapie brain homogenates. RML infection modestly but significantly upregulated *tnfα*, *il1β*, *ccl2* and *caspase-1* in WT, but not IKK KO, mixed glia. A significant decrease was seen *tnfα* (Fig 1C), *il1α* (Fig 1D), *il1β* (Fig 1E), and *ccl2* (Fig 1F) in both NBH-treated and RML-infected IKK KO cultures compared to RML-infected WT cultures (*p*<0.0001). No significant differences between WT and IKK KO cultures were observed for either NBH-treated or RML-infected groups for *ccl5* (Fig 1G) or *il6* (Fig 1H). A significant decrease was also seen in genes downstream of the NF-κB pathway, involved in the formation of the NLRP3 inflammasome. *Nlrp3* (Fig 1I) and *caspase-*1 (Fig 1J) were both downregulated in IKK KO cultures compared to RML-infected WT cultures (*p*<0.0001), but no significant differences were observed for *il18* (Fig 1K).

Together, significant genes regulated by NF-κB, and influential in the downstream NLRP3 pathway, are upregulated in a primary mixed glial population when exposed to RML-infected brain homogenate. When IKK is removed from microglia, expression of many NF-κB-related genes decreases drastically, independent of exposure to prions.

### Mice with IKK KO microglia present accelerated prion disease

Degeneration of neurons in the hippocampus, particularly in the CA1 region, is a prominent sign of prion diseases such as RML mouse-adapted scrapie (28, 29). Mice intracranially inoculated (RML) were assessed for behaviors associated with hippocampal function – the ability to build nests and to burrow. Historically, wild-type (WT) mice do not show impaired nest building until 18 weeks post-infection (wpi)(30), whereas IKK KO mice built deficient nests in as little as 13 wpi (Fig 2A). Similar findings were seen in burrowing, as IKK KO mice decreased in burrowing behavior as early as 12 wpi (Fig 2B), while WT mice historically do not show a decline in burrowing until 17 or 18 wpi (30). Although early clinical signs of prion disease began to manifest in WT mice at 13 or 14 wpi, these do not become significantly different from NBH controls until 21 wpi, and do not warrant euthanasia until 23 or 24 wpi (30). IKK KO mice began showing significant clinical signs beginning at 14 wpi, and signs were advanced enough to perform euthanasia for all animals by 18 wpi (Fig 2C). All WT mice succumbed to disease by 23 wpi, with an average of 157 days post-infection (dpi) +/- 7 days, whereas IKK KO mice succumbed to disease by 18 wpi, with an average of 123 dpi +/- 8 days (*p*=0.0002, Fig 2D).

**Fig 2.**
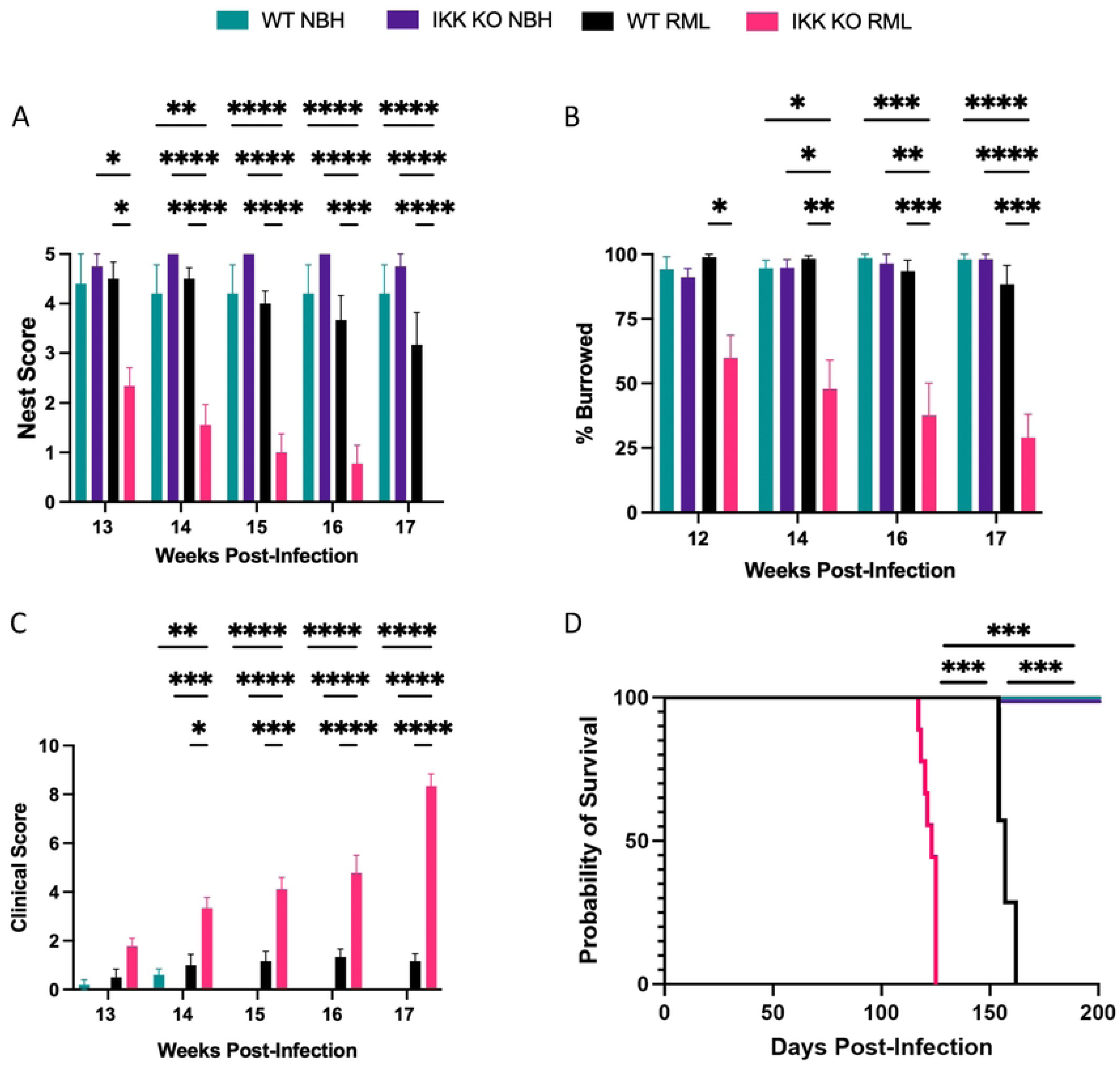
Mice with IKK KO microglia present accelerated prion disease. Mice were infected intracranially with RML mouse-adapted scrapie and monitored for changes in **A** nesting behavior, **B** burrowing behavior, and **C** clinical scores. **D** Animals displaying clinical scores of 10 were euthanized. NBH-treated groups contained 4 IKK KO and 5 WT mice, and RML-infected groups contained 9 IKK KO and 6 WT mice. One-way ANOVA and post-hoc Tukey test, error bars = SEM. For survival curve, a Log-rank (Mantel-Cox) test was performed. \**p* < 0.05, \*\**p* < 0.01, \*\*\**p*< 0.001, \*\*\*\**p* < 0.0001.

### Removal of IKK in microglia induces change in microglia number and morphology during prion infection

Although generally considered to provide protection in the host, the role of microglia in prion pathogenesis remains poorly understood (31). Mice with IKK KO microglia succumbed to prion infection around 17 wpi. To understand the pathological processes that contributed to this rapid disease onset in mice with IKK KO microglia, a cohort of prion-infected WT mice was euthanized at 17 wpi for comparison, referred to as age-matched WT mice. Comparisons were made between terminal mice with IKK KO microglia and both age-matched WT and terminal WT mice. The number of Iba1+ microglia is known to increase over the course of prion infection (32, 33). Brain regions associated with significant prion deposition in RML scrapie were assessed for Iba1+ microglia in the frontal cortex, hippocampus, thalamus and cerebellum of prion-infected WT and IKK KO mice (Fig 3A)(32, 34). Significantly more Iba1+ microglia were detected in the cortex (*p*=0.0424, Fig 3B), hippocampus (*p*=0.0130, Fig 3C), and thalamus (*p*=0.0096, Fig 3D), but not the cerebellum (p=0.0722, Fig 3E), in mice with IKK KO microglia. No significant differences were seen in Iba1+ microglia between the two groups when they were mock-infected (Supplemental Fig 2).

**Fig 3.**
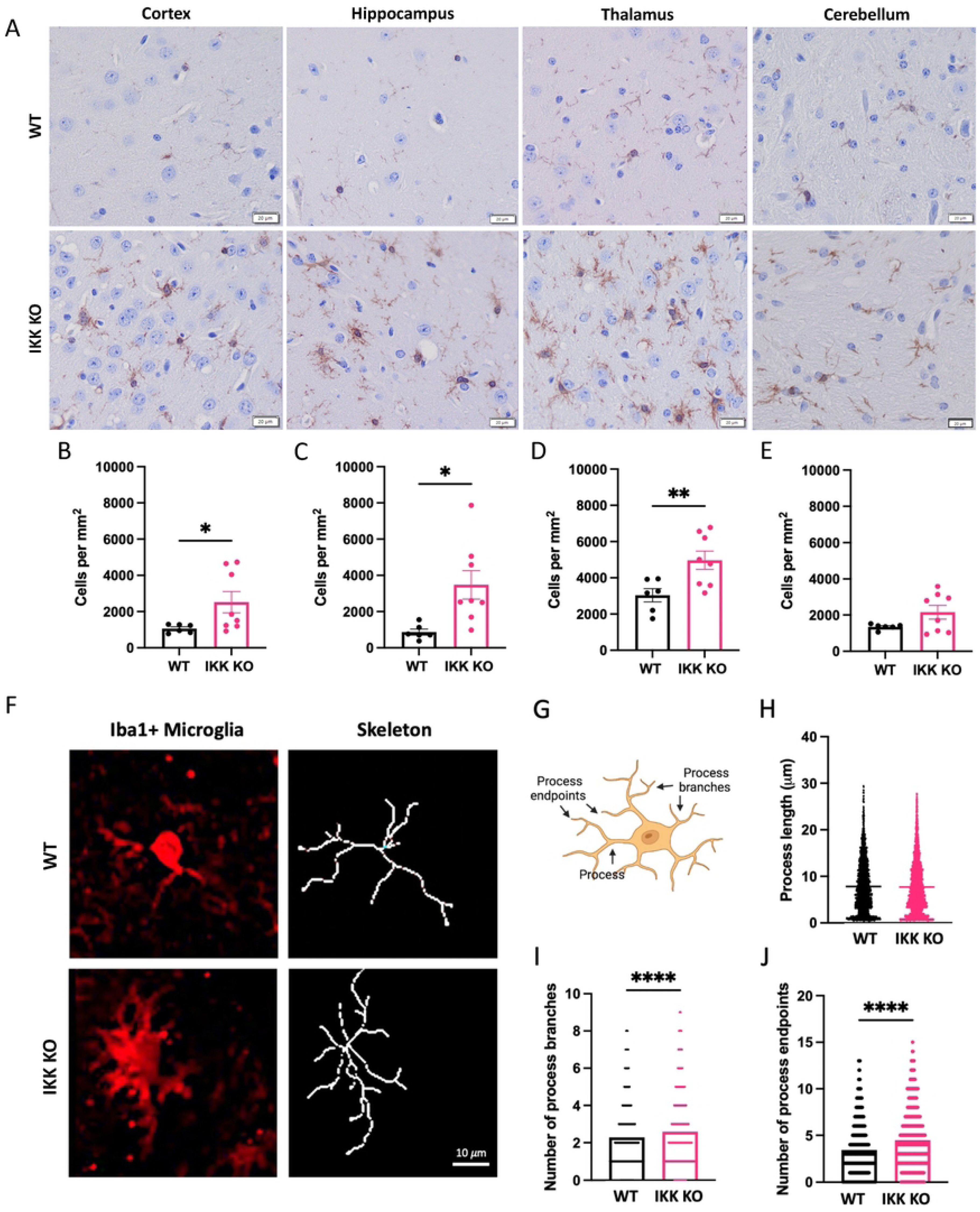
Removal of IKK in microglia induces changes in microglia number and morphology during prion infection. Terminal mice with IKK KO microglia were compared to age-matched WT infected mice. **A** Brains were stained for Iba1+ microglia, which were counted and compared in the **B** cortex, **C** hippocampus, **D** thalamus and **E** cerebellum. **F** Representative images of Iba1+ hippocampal microglia morphological skeletons between groups. **G** Microglia cartoon depicting features analyzed via skeletonization. **H** Process length, **I** number of process branches and **J** number of process endpoints was compared between hippocampal microglia from WT mice and those with IKK KO microglia. Welch’s t-test, error bars = SEM, \**p* < 0.05, \*\**p*< 0.01, \*\*\*\**p* < 0.0001.

The morphology of Iba1+ hippocampal microglia was analyzed using skeletonization (Fig 3F) to assess process length, the number of process branches, and the number of process endpoints for each cell (Fig 3G)(30). Although process length was not significantly different between groups (Fig 3H), the number of process branches (Fig 3I) and process endpoints (Fig 3J) were significantly greater in mice with IKK KO microglia (*p*<0.0001).

Together, these data show that mice with IKK KO microglia still have equivalent numbers of Iba1+ microglia in the brain. However, when intracranially infected with RML scrapie, these mice showed a rapid increase in the number of Iba1+ microglia in the cortex, hippocampus and thalamus compared to age-matched infected WT mice. Analysis of the morphology of hippocampal microglia showed more activation in IKK KO microglia compared to those from WT mice.

### Brains with microglial IKK KO show increased GFAP expression and activated astrocytes during prion infection

GFAP+ C3+ (activated) astrocytes are known to increase over the course of prion infection (35, 36). GFAP+ astrocytes were counted in the frontal cortex, hippocampus, thalamus and cerebellum of prion-infected WT and IKK KO mice (Fig 4A). Significantly more GFAP+ astrocytes were detected in the cortex (*p*=0.0020, Fig 4B), hippocampus (*p*<0.0008, Fig 4C), thalamus (*p*<0.0037, Fig 4D) and cerebellum (*p*=0.0107, Fig 4E) in mice with IKK KO microglia. No significant changes were seen in GFAP+ astrocyte numbers in mock-infected brains, except for the cortex where more cortical GFAP+ astrocytes were present in mice with IKK KO microglia compared to WT mice (p=0.0027, Supplemental Fig 3).

**Fig 4.**
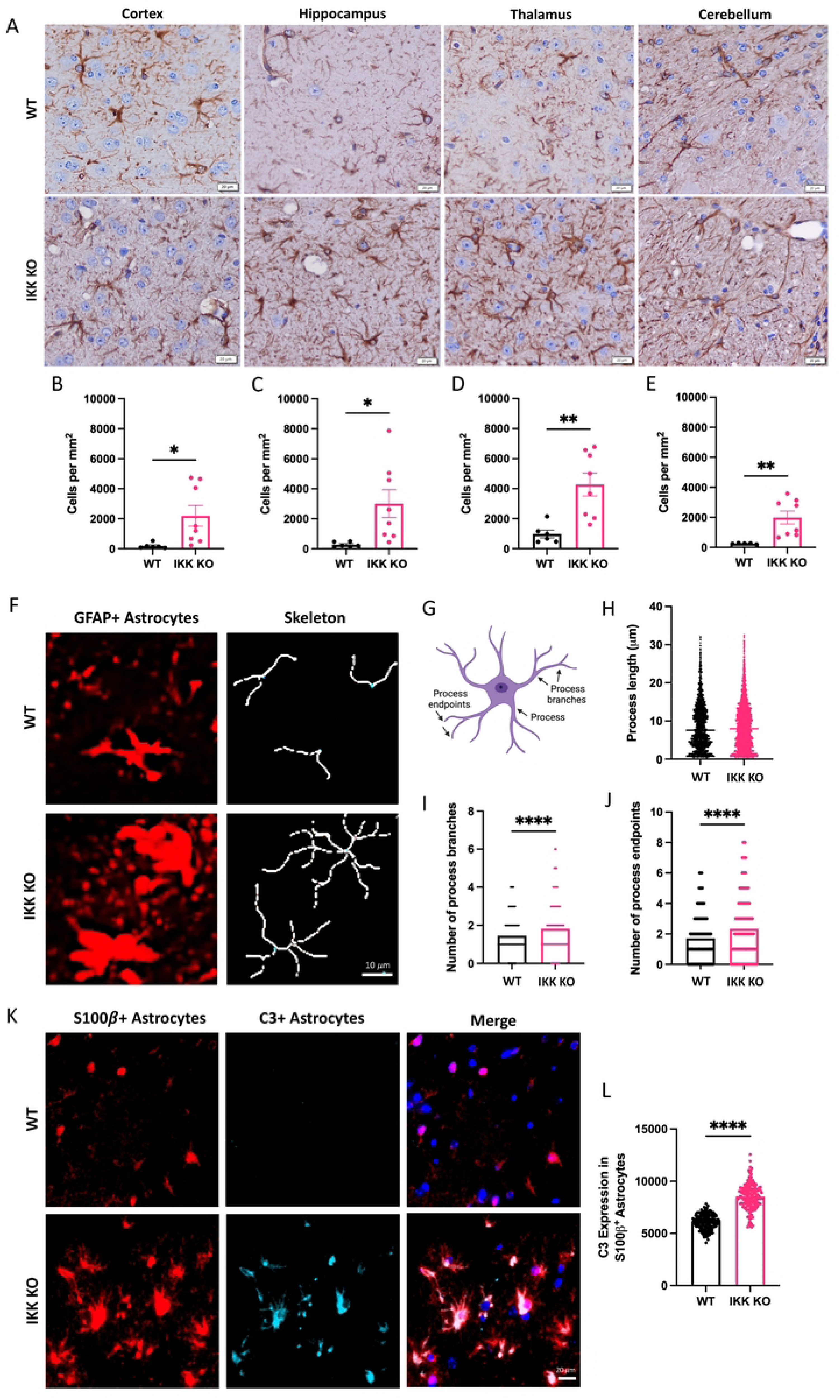
Brains with microglial IKK KO have increased GFAP expression and activated astrocytes during prion infection. Terminal mice with IKK KO microglia were compared to age-matched WT infected mice. **A** Brains were stained for GFAP+ astrocytes, which were counted and compared in the **B** cortex, **C** hippocampus, **D** thalamus and **E** cerebellum. **F** Representative images of GFAP+ hippocampal astrocyte morphological skeletons between groups. **G** Astrocyte cartoon depicting features analyzed via skeletonization. **H** Process length, **I** number of process branches and **J** number of process endpoints was compared between hippocampal astrocytes from WT mice and those with IKK KO microglia. **K** Hippocampal astrocytes were co-stained for the pan-astrocytic marker S100β and the reactive astrocyte marker C3. **L** Mean grey intensity of C3 was compared in S100β+ astrocytes (arbitrary units). Welch’s t-test, error bars = SEM, \**p* < 0.05, \*\**p*<0.01, \*\*\*\**p* < 0.0001.

The morphology of GFAP-expressing hippocampal astrocytes was analyzed using skeletonization (Fig 4F) to assess process length, the number of process branches, and the number of process endpoints for each cell (Fig 4G)(30). Although process length was not significantly different between groups (Fig 4H), the number of process branches (Fig 4I) and process endpoints (Fig 4J) were significantly greater in mice with IKK KO microglia (*p*<0.0001).

To assess the number of neurotoxic reactive astrocytes present in the hippocampus (5, 14), brains were stained for colocalization of the pan-astrocyte marker S100β and the complement cascade protein C3 (Fig 4K). Significantly more C3 was detected in S100β+ hippocampal astrocytes in mice with IKK KO microglia (*p*<0.0001, Fig 4L).

Healthy mice with IKK KO microglia have similar numbers of GFAP+ astrocytes in most brain regions analyzed. Infected mice with IKK KO microglia showed a rapid increase in the number of GFAP+ astrocytes in the cortex, hippocampus, thalamus and cerebellum compared to age-matched WT mice.

### Removal of IKK in microglia does not protect against prion-induced neuronal death *in vitro* or *in vivo*

To determine the effects of factors secreted into the media by wild-type (WT) mixed glia compared to mixed glia with IKK KO microglia, glial conditioned media (GCM) was harvested from NBH or RML-treated primary mixed glial cultures after a 96-hour incubation. N2a neuroblastoma cells were incubated in GCM for 48 hours, then viability was measured with Presto Blue (Fig 5A). GCM from RML-infected WT glia had an average of 13.4% fewer viable cells (*p*<0.01), and RML-infected IKK KO glia had an average of 23.9% fewer viable cells (*p*<0.0001), compared to the NBH-treated samples (Fig 5B). However, there was no statistically significant differences in neuronal viability between WT and IKK KO RML-infected cell GCM.

**Fig 5.**
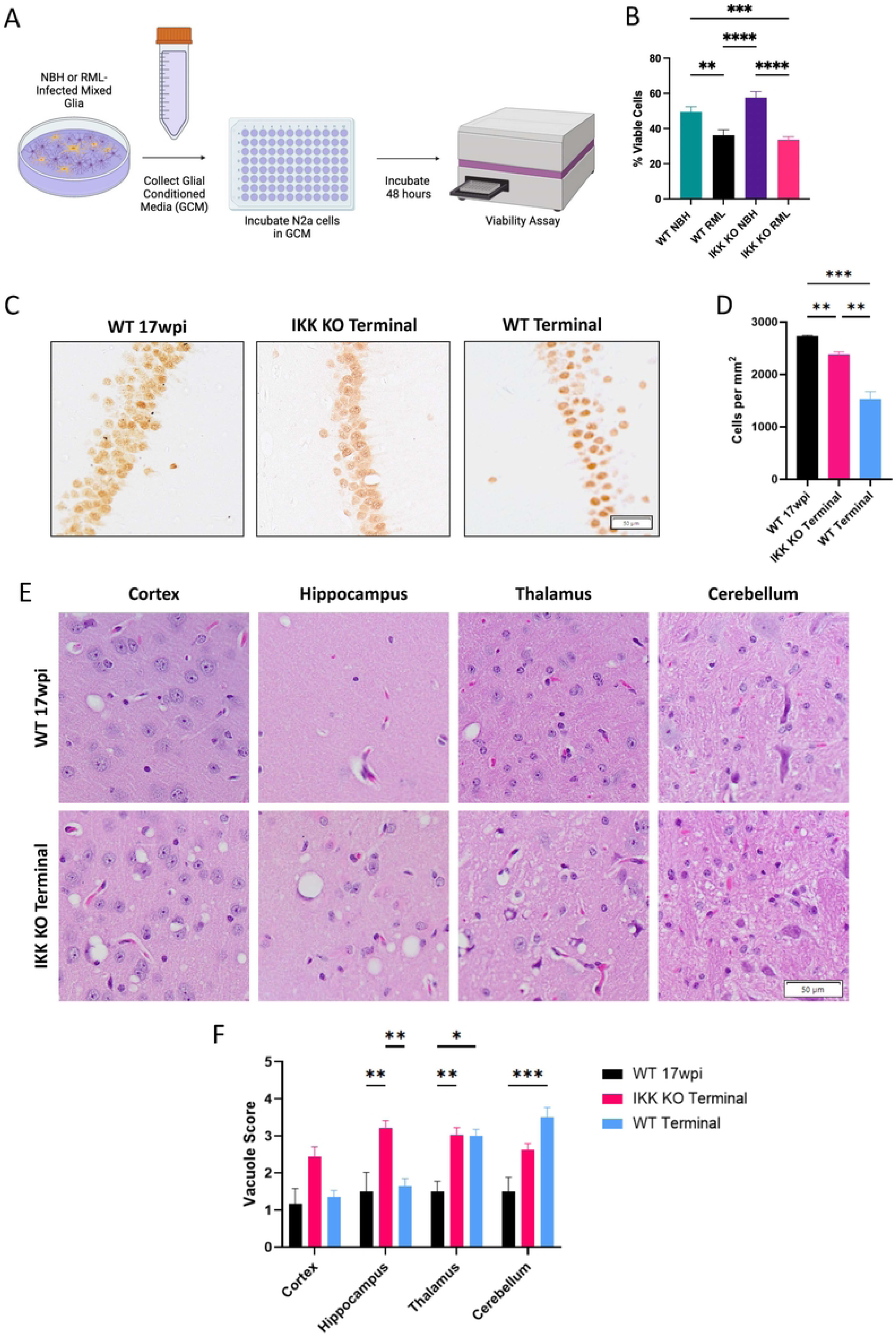
Removal of IKK in microglia does not protect against prion-induced neuronal death *in vitro* or *in vivo.* **A** Mixed glial cultures were treated for 7 days with NBH or RML homogenates. Glial conditioned media (GCM) was isolated and plated on N2as in 96-well plates and chamber slides for 48 hours. **B** GCM-treated N2as were analyzed for percent viability using a Presto Blue cell viability assay. **C** Representative images of NeuN+ cells in the CA1 region of the hippocampus for terminal mice with IKK KO microglia, infected age-matched WT mice and terminal WT mice and **D** CA1 NeuN+ neuronal counts. The severity of vacuolization for infected mice was scored from 1-5 by three blinded pathologists in the cortex, hippocampus, thalamus and cerebellum. **E** Representative images of vacuoles for terminal mice with IKK KO microglia and infected age-matched WT mice. **F** Bar graph showing mean vacuole severity for all infected mice. For the viability assay, a one-way ANOVA and post-hoc Tukey test was used. For NeuN counts, a Brown-Forsythe and Welch ANOVA was used. For vacuole counts, a two-way ANOVA and post-hoc Tukey test was used. Error bars = SEM, \**p* < 0.05, \*\**p* < 0.01, *** *p* < 0.001, \*\*\*\**p* < 0.0001.

Neuronal loss in prion-infected mice predominantly occurs in the CA1 region of the hippocampus (28). To assess the number of living neurons, NeuN+ neurons were counted in the CA1 region to assess for neuronal loss. No differences were observed between the CA1 regions of hippocampi from mock-infected mice with WT and IKK KO microglia (Supplemental Fig 4A). Significantly fewer NeuN+ neurons were detected in mice with IKK KO microglia, indicating significant hippocampal neuronal loss compared to infected age-matched WT mice (*p*=0.0015, Fig 5C-D). Terminal WT mice had significantly fewer NeuN+ neurons compared to both terminal mice with IKK KO microglia and infected age-matched WT mice (*p=* 0.0014 and *p*= 0.0004, respectively).

A pathological hallmark of prion disease is the development of a spongiform pattern within the brain, referred to as vacuoles (37, 38). Brains were stained with hematoxylin and eosin (H&E) to assess for severity of vacuoles based on both size and number in the frontal cortex, hippocampus, thalamus and cerebellum (Fig 5E). Vacuole severity was significantly higher in the hippocampus of infected mice with IKK KO microglia compared to infected age-matched and terminal WT mice (*p=*0.0015 and *p*=0.0029, respectively), and higher in the thalamus compared to age-matched WT mice (*p=*0.0079, Fig 5F). No significant vacuoles were seen in any of the above brain regions for mock-infected mice (Supplemental Fig 4B-E).

Infected mice with IKK KO microglia presented with signs of neurodegeneration and vacuolization at an accelerated pace compared to age-matched WT mice. The cell viability assay shows that primary glia from both WT and IKK KO mice are secreting neurotoxic factors, even in the absence of functional IKK in the microglia.

### IKK KO in microglia leads to increased accumulation of PK-resistant PrP *in vitro* but decreased accumulation *in vivo*

Cell lysates from infected primary mixed glia containing WT astrocytes and WT or IKK KO microglia were analyzed for infectious, PK-resistant PrP (denoted PrP^Sc^) or total PrP (containing both PrP^C^ and PrP^Sc^) via western blot (Fig 6A). Comparison of band densitometry revealed a significant increase in PrP^Sc^ (*p*<0.0001, Fig 6B) and total PrP (*p*<0.001, Fig 6C) from lysates with IKK KO microglia compared to those with WT microglia.

**Fig 6.**
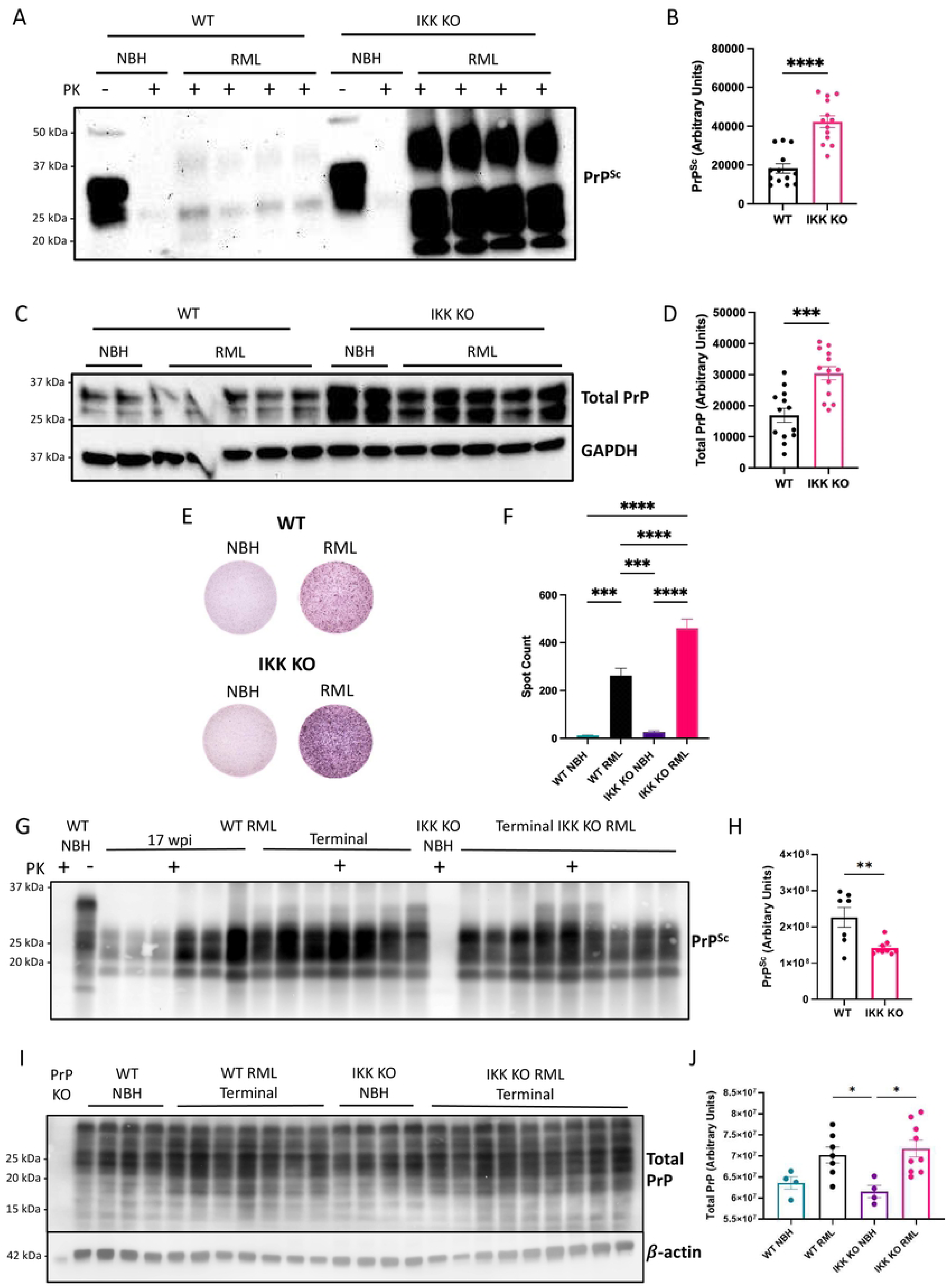
IKK KO in microglia leads to increased accumulation of PK-resistant PrP *in vitro* but decreased accumulation *in vivo*. Mixed glial cells containing WT astrocytes and WT or IKK KO microglia were treated with NBH or infected with RML for 7 days. **A** Cells were lysed and digested with PK for PrP^Sc^ observation and **B** protease-resistant protein signal was measured. **C** Undigested protein was assessed for total PrP and **D** protein signal for total PrP was measured. **E** Infected cells were trypsinized and transferred to an ELISpot plate for a scrapie cell assay to **F** count the number of infected cells (normalized). Prion western blot analysis and spot counts are combined data from three separate experiments. **G** Homogenized brains from terminally infected IKK KO mice and both age-matched and terminally infected WT mice were digested with PK and **H** protease-resistant protein signal was measured using Sha31 antibody. **I** Total PrP was assessed using Sha31 antibody for both terminally infected and mock-infected mice and **J** compared between groups. For western blot quantification of PrP^Sc^, an unpaired t-test was performed. For scrapie cell assay and western blot analysis of total PrP, a one-way ANOVA and post-hoc Tukey test was used. Error bars = SEM, \**p* < 0.05, \*\**p* < 0.01, *** *p* < 0.001, \*\*\*\**p* < 0.0001.

Infected mixed glia were quantified for prion plaques using a scrapie cell assay (Fig 6D). Significantly more plaques (spots) were detected in RML-infected WT cells compared to the NBH-treated WT cells (*p*<0.001) and in RML-infected IKK KO cells compared to NBH-treated IKK KO cells (*p*<0.0001). Similar to the western blot analyses, there were significantly more prion plaques present in RML-infected cultures containing IKK KO microglia compared to WT microglia (*p*<0.0001) (Fig 6E).

To determine if this phenomenon was retained *in vivo*, terminal IKK KO animals and both age-matched and terminal WT animals were analyzed for prion accumulation in the brain. The left hemisphere was homogenized and assessed for PK-resistant PrP by immunoblot. Multiple antibodies were used to probe for PrP, including an N-terminal antibody (12B2) and a C-terminal antibody (Sha31) (antibody epitopes shown in Supplemental Fig 5A). No differences were seen in band densitometry or glycosylation pattern between infected terminal IKK KO and age-matched WT brains using either 12B2 (Supplemental Fig 5B-C) or Sha31 (Supplemental Fig 6A-B). No changes were seen in total PrP in these brains with Sha31 (Supplemental Fig 6C). At this time point there were no detectable differences in total PrP between NBH control and RML infected WT brains, whereas there was significantly more total PrP in RML-infected IKK KO brains compared to NBH control IKK KO brains (p<0.001, Supplemental Fig 6D).

The C-terminal antibody, Sha31, was used to analyze prion accumulation comparing terminal WT mice and terminal IKK KO mice. Significantly more PK-resistant PrP was detected in terminal WT mice compared to terminal IKK KO mice (p<0.0045, Fig 6G-H). Total PrP was the same between terminal WT and IKK KO mice (Fig 6I-J). Assuredly, brains from mice with IKK KO microglia showed a decrease in IKKβ of approximately 50% compared to WT brains at terminal stages, which was independent of infection (Supplemental Fig 7).

## Discussion

The exact neurotoxic agent in prion disease is not fully understood, as expression of PrP is critical for cells to become infected and for neuronal loss (31, 39-41), but glial cells also play a distinct role in disease pathogenesis (36, 42, 43). Neurons and astrocytes express the highest amount of PrP in the body and brain (44), and therefore are easily infected with PrP^Sc^ and can traffic it between cells (45). Microglia scavenge and phagocytose PrP^Sc^, which results in a phenotypic change from scavenging and ramified to activated or amoeboid, often denoted as a shift from M2 to M1, or homeostatic microglia to disease-associated microglia (DAM)(6, 31, 33, 34, 42, 46). This induces the secretion of factors such as pro-inflammatory cytokines and chemokines, which in turn can induce astrocytes to become reactive and produce inflammatory and neurotoxic mediators (5, 14, 27).

Animal models of prion disease which have a reduction or elimination of microglia have varying effects on disease outcome (34, 47-51), depending on the time point in which the reduction occurs. Generally, microglia are found to be protective to the host, and disease worsens when microglia are decreased or removed. Similarly, genetic manipulation to change the inflammatory state of astrocytes have not successfully extended the lives of mouse models (5, 51). Together, these findings suggest that inflammation from glial cells plays a critical role in host protection but can become detrimental if left unchecked. Therefore, reduction, but not elimination, of glial-induced inflammation, may be a promising avenue for therapeutics (24, 52-54). To best develop treatments, further investigation is required to understand the involvement of specific inflammatory signaling pathways from astrocytes and microglia, and how they affect neuronal health in the prion-infected brain.

Microglia, potent regulators of prion infection, are involved in host protection through inflammatory signaling, communication with astrocytes, phagocytosis and degradation of PrP^Sc^ (31, 47, 49-51, 55). Many genes involved in the NF-κB signaling pathway are highly upregulated in animal models with prion disease (4, 8, 9), suggesting that this pathway is a critical innate immune response to prion infection. To date, no studies have analyzed the role of microglia-specific NF-κB signaling in a mouse model. One study removed IKK from cells in the CNS with neuroectodermal lineage – namely neurons, oligodendrocytes, and astrocytes. No changes in disease time-course, PrP^Sc^ accumulation or astrogliosis were found and the authors concluded that NF-κB does not play a significant role in prion-induced inflammatory signaling (13). Another study directly knocked out the subunits of NF-κB in cells derived from the neuroectoderm. This group saw a decrease in survival in these mice upon prion inoculation, and increased apoptosis (12). Both of these studies failed to account for microglia, which are derived from the mesoderm (56) and known to be critical regulators of NF-κB-associated inflammation in prion and other diseases (5, 14, 27, 51). Here, we utilize both a primary cell model that contains wild-type (WT) astrocytes and IKK KO microglia (23), as well as a mouse model with IKK KO microglia, to characterize the role of microglia-specific IKK and NF-κB signaling in prion disease.

Cx3Cr1Cre mice express Cre recombinase under the *cx3cr1* promoter in the mononuclear phagocyte system and were combined with floxed IKKβ mice (23) to generate mice with IKK knockout (KO) specific to mononuclear phagocytes (macrophages, monocytes and microglia). As macrophages and monocytes are rarely found in the brain in prion disease (27), these mice are effectively microglia-specific IKK KO. In an innate immune response, IKK is critical for NF-κB translocation to the nucleus for transcription (19, 20), making these cells ineffective at NF-κB signaling. However, importantly, all other brain cells in these mice have functional IKK and NF-κB signaling. Analysis of brain homogenates by western blot reveal a 50% decrease in IKKβ in IKK KO mice compared to WT brains (Supplemental Fig 7).

It has been established that even a small number of microglia can greatly impact a cell culture system, especially in modulation of astrocytes (57). There is significant signaling cross-talk between astrocytes and microglia, as astrocytes have been shown to remain relatively unresponsive to environment changes in the absence of microglia-derived signaling (14, 17), and microglia rely heavily on astrocytes to promote their proliferation (25). To better elucidate signaling between these cell types, we infected primary mixed glia containing WT astrocytes and WT or IKK KO microglia with prions. Astrocytes highly express CSF1, which - alongside IL34, is expressed by neurons - binds to colony stimulating factor 1 receptor (CSF1R) on microglia to promote their proliferation and survival (25). Interestingly, one of the few upregulated genes in the infected mixed glial cultures containing IKK KO microglia was CSF1 (Fig 1), suggesting that astrocytes in these cultures are producing increased CSF1 to promote microglial survival. CSF1 is highly upregulated in the brain in response to inflammatory-inducing signals such as lipopolysaccharide (LPS) (58).

Nearly 70% of the 84 NF-κB-associated genes analyzed in the primary mixed glia cultures were downregulated in RML-scrapie-infected IKK KO glia compared to RML-infected WT glia (Fig 1, Supplemental Table 1 for complete list). Of these genes, many were inflammatory cytokines and chemokines known to be upregulated in the prion-infected brain (8, 9, 27). Some of these cytokines, including tnfα, il1α and il1β, are predominantly produced by microglia and influence the state of astrocyte reactivity (5, 14). CCL2 is produced by astrocytes and promotes an M1 phenotype in microglia (59). This suggests that in the absence of microglia-specific NF-κB signaling, signaling from astrocytes is also decreased, as microglia are critical for astrocytes to reach a full activation state (60). These results highlight that crosstalk and reciprocal activation of microglia and astrocytes are important for proper function of both cell types in response to infection, and the role of microglia-specific NF-κB signaling in this activation.

To further interrogate the influence of microglia-specific IKK KO on specific genes of interest, we performed qPCR analysis on three to five sets of RML-infected primary mixed glia, each isolated from a separate set of mouse pups (Fig 1). *Tnfα*, *il1α* and *il1β* were consistently downregulated in IKK KO glia, as were the NLRP3-associated genes *nlrp3* and *caspase-1* suggesting that these genes are predominantly upregulated by NF-κB within microglia. We also saw an overall downregulation in *ccl2*, although less robust than the other genes. As this chemokine is produced mainly by astrocytes (59), it is likely still expressed in the absence of microglial NF-κB signaling. Due to known involvement in NF-κB signaling and prion pathogenesis, we additionally looked at expression of *ccl5* and *il6*, as well as the NLRP3-associated gene, *il18*. A large variation in expression of these genes with each set of glia analyzed showed both significant downregulation and, intriguingly, significant upregulation, suggesting compensatory signaling pathways from astrocytes, microglia, or both, that contribute to neuroinflammatory response. However, unlike many of the other genes, no significant changes were seen between the WT glia treated with normal brain homogenate (NBH) or RML, suggesting that this inflammation is not directly induced by prions.

Genes significantly downregulated in these cells include toll-like receptors (TLRs), particularly *tlr1*, *tlr2*, *tlr4*, *tlr6* and *tlr9*, all of which are highly expressed in microglia. Decrease in TLR expression is consistent with a decrease in microglial numbers, and also likely contributes to an overall decrease in inflammatory molecules (58), as signaling is dysregulated in the absence of sufficient TLRs. *Tlr3* is slightly downregulated, but this TLR is highly expressed in astrocytes, along with low-level expression of *tlr1*, *tlr4*, *tlr5* and *tlr9* (61). Interestingly, microglia are critical for TLR4 signaling in astrocytes, and important for optimal signaling through TLR2 and TLR3 (62), which may contribute to overall downregulation of astrocyte-specific signaling in cultures containing IKK KO microglia, as these microglia cannot fully prime the astrocytes. TLR4 is one of the best studied TLRs, as its stimulation by signals such as LPS induce NF-κB inflammation in microglia (20, 58), which is critical to elicit an astrocyte response (62). In prion mouse models, *tlr1*-*9* are all upregulated in the brain at terminal stages of disease (58). TLR2 and TLR4 signaling are important for sensing damage-associated molecular patterns (DAMPs) and has been shown to be implicated in prion disease, as knocking out either of these receptors in mice accelerates disease (63, 64).

Despite KO microglia lacking IKK2 and therefore having non-functional canonical NF-κB signaling, the Iba1+ microglia in infected IKK KO mice increase in numbers more rapidly than those in age-matched infected WT mice. Significantly more Iba1+ microglia were observed in most of the analyzed brain regions, most of which were characteristic of M1 microglia or DAM, having a large amoeboid shape with increased processes and process branches, whereas those from WT mice still resembled an M2 phenotype, with fewer processes and a smaller cell body (Fig 3). This suggests that microglia are still reacting to the prion-infected brain even in the absence of IKK and NF-κB signaling, and that their activation state may be dysregulated. Further interrogation is required to determine if this activated phenotype is a direct or indirect response to IKK knock-out.

GFAP expression increased in all brain regions analyzed in terminally infected mice with IKK KO microglia compared to age-matched infected WT mice. RML-infected mice with IKK KO microglia had more reactive astrocytes in the hippocampus, demonstrated by increased C3 expression and morphology (5, 14)(Fig 4). This suggests that NF-κB signaling from microglia may be a critical regulator of astrocyte activation states in prion pathogenesis, and astrocytes may be compensating in response to dysfunctional microglia (51).

Studies demonstrate that decreasing or fully ablating microglia in prion-infected animals leads to accelerated disease, increased vacuolization and astrogliosis (47, 51). Astrocytes from microglia-ablated mice showed enhanced ability to phagocytose neuronal synapses and increased unfolded protein response, both of which contribute to irreversible neuronal loss (29, 51). TNFα and IL1α are decreased in prion-infected mice without microglia, suggesting that microglia are a key source of NF-κB-related proinflammatory cytokines (47). These microglia-derived cytokines, alongside C1qa, are responsible for inducing astrocyte reactivity, marked by increased C3 expression (14). Increased inflammatory signaling by microglia is consistent with its role as the main antigen presenting cell with associated helper cell-like functions in the brain. Knockout of TNFα, IL1α and C1qa in prion-infected mice, although sufficient in ablating C3+ astrocytes, also decreased survival time and the number of homeostatic microglia (5). Together, these studies show that microglia and astrocytes tightly regulate one another in the prion-infected brain and unbalancing this relationship leads to dysfunction and accelerated disease.

Microglial-induced inflammation is often cited as an inducer of neuronal cell death, either directly or indirectly (5, 14, 17, 46, 51, 65, 66). Therefore, it was surprising that glial conditioned media (GCM) from infected WT and IKK KO mixed glial cultures showed similar neurotoxicity to N2a cells in cell viability assays (Fig 5A-B). This suggests that even in the absence of NF-κB signaling from microglia, glial cells are secreting neurotoxic factors similar to WT cells. However, these findings were consistent with our *in vivo* data, which showed neuronal death both directly and indirectly in animals with IKK KO microglia. Significantly fewer NeuN+ neurons were observed in the CA1 region of the hippocampi in terminally infected mice with IKK KO microglia compared to age-matched infected WT mice. This was consistent with the decrease in burrowing and nesting behavior, both behaviors that are related to neuronal health in the hippocampus. Terminal WT mice had significantly fewer NeuN+ neurons compared to terminal mice with IKK KO microglia (Fig 5C-D). There is evidence that vacuole formation is an indirect marker of neuronal death (38, 67), and terminally infected IKK KO mice showed significantly more vacuolization in the hippocampus than either age-matched or terminally infected WT mice (Fig 5E-I). This finding is consistent with a recent study that described increased vacuoles but no changes in CA1 hippocampal neurons in prion-infected mice lacking microglia (51). Together, our findings suggest that IKK and NFκB signaling from hippocampal microglia are critical for regulating astrocytes and protecting hippocampal neurons (51). Further interrogation is required to understand this complex cellular crosstalk between these cell types.

Cellular responses to misfolded proteins are impaired in the primary cell cultures containing IKK KO microglia, demonstrated by increased accumulation of PrP^Sc^ via western blots and scrapie cell assays (Fig 6A-E). Intriguingly, this phenomenon is not observed when looking at crude brain homogenates from infected mice. Brains from terminally infected mice with IKK KO microglia show no difference in either PK-resistant PrP or total PrP compared to age-matched infected WT mice (Supplemental Fig 6), but decreased amounts of PK-resistant PrP and total PrP compared to terminal WT mice (Fig 6). This suggests that mice with IKK KO microglia succumbed to disease quicker than their brains accumulate misfolded prions, suggesting that neuronal death is induced by factors besides PrP^Sc^, such as abnormalities in glia. This phenomenon has been described for other mouse models. Mice with total microglia ablation succumb to disease due to astrocyte dysfunction, despite having less PrP^Sc^ compared to controls (51).

Accumulation of misfolded proteins is commonly associated with dysregulation of autophagy and lysosomal function. Rescue of either of these pathways is shown to improve life expectancy and clinical pathology in prion-infected mice (68-70). Misfolded proteins are first identified and ubiquitinated by chaperone proteins, leading to the binding and sequestration of aggregates by p62, a protein upregulated by the NF-κB pathway (71, 72). Moreover, signals that activate the IKK complex are shown to initiate autophagy even in the absence of downstream NF-κB signaling, suggesting that knockout of IKK leads to impaired autophagy regardless of NF-κB-induced p62 signaling (22). Increased abnormal protein accumulation was demonstrated in mice with IKK KO microglia in a model of rotenone-induced Parkinson’s disease (PD). However, conversely to our findings, there was a significant reduction in reactive astrocytes and neuronal loss in rotenone-treated male mice with microglial IKK KO, as well as a decrease in clinical signs of PD (3). Studies have demonstrated that microglia are the predominant source of NF-κB signaling in animal models of neurodegeneration such as amyotrophic lateral sclerosis and tauopathy, and inhibition of this pathway in microglia in these models proves neuroprotective (1, 2). Knocking out IKK2/IKKβ in microglia in a mouse model of tauopathy led to increased accumulation of tau in microglia, but prevented its seeding and further spreading, ultimately leading to neuroprotection (1). These studies indicate IKK as a critical mediator of aggregated protein modification and clearance. Our findings suggest that increased prion accumulation associated with IKK KO in microglia does not increase levels of total brain PrP^Sc^, and may be limited to astrocytes and microglia, or limited to our primary cell culture model. Further studies are underway to tease out the mechanism by which these glial cells are responsible for enhanced prion accumulation.

Further characterization of this model is necessary to understand how microglia-specific IKK KO affects NF-κB signaling, as well as downstream effects such as the involvement of other inflammatory pathways, autophagy, and neuronal cell death. Although we see profound changes in our cell culture and mouse model in response to prion infection, it is difficult to tease out whether this is due to the role IKK plays directly on protein clearance in glial cells, the lack of sufficient NF-κB inflammatory signaling, or a combination of the two. More intriguingly, we do not see increased accumulation of PK-resistant prions in the brains of infected mice with IKK KO microglia. These mice show similar amounts of PrP^Sc^ compared to age-matched WT mice, and less PrP^Sc^ compared to terminal WT mice. However, neurons are rapidly deteriorating, and infection is fatal in mice with IKK KO microglia at a much earlier timepoint suggesting that there are other factors contributing to neurotoxicity other than prion accumulation – likely due to dysfunctional microglia and their crosstalk with astrocytes. Future studies will focus on disease progression by analyzing glial and neural morphology and function, prion propagation and clearance in prion-infected IKK KO mice prior to their display of clinical signs. This model gives insight to an age-old question – is it prion aggregation and accumulation, or downstream effects such as inflammation and gliosis, that leads to neurodegeneration? Understanding the involvement of IKK and NF-κB signaling in microglia may uncover both biomarkers and therapeutic targets for prion diseases.

## Methods

### Animal care and ethics

Cx3Cr1Cre-IKKflox mice (C57Bl6/J background, Cx3CrlCre available from Jax #025524) were kindly provided to us by Dr. Ronald Tjalkens (3). All mice were bred and maintained at Lab Animal Resources, accredited by the Association for Assessment and Accreditation of Lab Animal Care International. All protocols are in approved by the Institutional Animal Care and Use Committee at Colorado State University. An approximately equal number of male and female mice were used throughout the study.

### Isolation of mixed glia

Zero to two-day old C57Bl6/J or CX3CR1-IKKflox pups were euthanized and brains were extracted. Cerebellum, midbrain and hippocampus were removed and discarded, and the brains were placed in MEM/EBSS (Hyclone) containing 2x penicillin/streptomycin/neomycin (PSN) (Sigma) on ice. Cortical tissue was dissociated using 1.5 U/ml Dispase to isolate mixed glial cultures, as described previously (24). 10^6^ mixed glial cells were plate in 10 cm dishes and cells were incubated at 37 ° C with 5% CO_2_ in glial growth media (MEM/EBSS + 10% FBS + 1% PSN). Media was replaced after 24 hours and changed weekly.

### Infection of primary glial cell cultures

For *in vitro* prion infection, mixed glia were plated at 10^5^ cells per well in 6-well plates and infected at 80% confluency with 0.1% normal or RML brain homogenate in media. Media was removed 72 hours later and cells were washed twice with PBS prior to fresh media being added to remove any residual brain homogenate. Supernatants (glial conditioned media (GCM)) and protein or RNA were extracted from plates after an additional 96 hours (7 days from initial homogenate treatment) and analyzed as described below.

#### Reverse transcriptase quantitative PCR analysis

RNA was extracted from 6-well dishes using cell scraping, QIAshredder and RNeasy extraction kits, in accordance with manufacturer’s protocol, including a DNase digestion step with the RNase free DNase kit (Qiagen, Valencia, CA). Purity and concentration were determined using a ND-1000 spectrophotometer (NanoDrop Technologies, Wilmington, DE). Following isolation and purification, 25 ng per sample of RNA was reverse transcribed using the iScript Reverse Transcriptase kit (BioRad, Hercules CA). cDNA was amplified within 24 hours of reverse transcription using iQ SYBR Green Supermix (BioRad, Hercules CA). For NF-κB panel, the RT^2^ profiler PCR array for Mouse NF-κB Signaling Pathway (Qiagen) was used with Iq Sybr (Bio-Rad), following manufacturer’s protocol. Plates were analyzed using the Bio-Rad CFX96 Real-Time System. Files were uploaded into RT^2^ profiler PCR array analysis software for quality control and statistical analysis. Samples were normalized to 5 reference genes selected by the analysis software – Hsp90ab1, Tollip, Irak1, Bcl2l1 and Tnfrsf1a. For qPCR of specific genes, the corresponding validated primer sequences were used for each gene at 10mM. Plates were analyzed using the LightCycler 480 II (Roche). The expression data was analyzed using the 2-ΔΔCT method and normalized to expression of reference genes *β-actin.* The fold difference was compared to control (WT normal brain homogenate treated) samples (73). All treatments were done in triplicate and cDNA was measured in triplicate. Data is a combination of three to five biological replicates. Validated primer sequences are listed in Table 1.

**Table 1.**
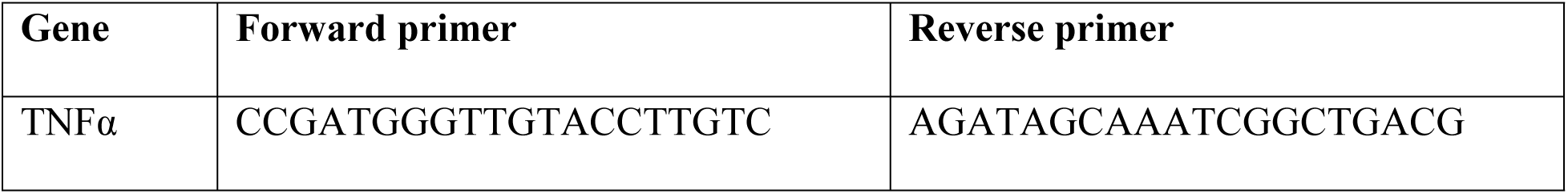
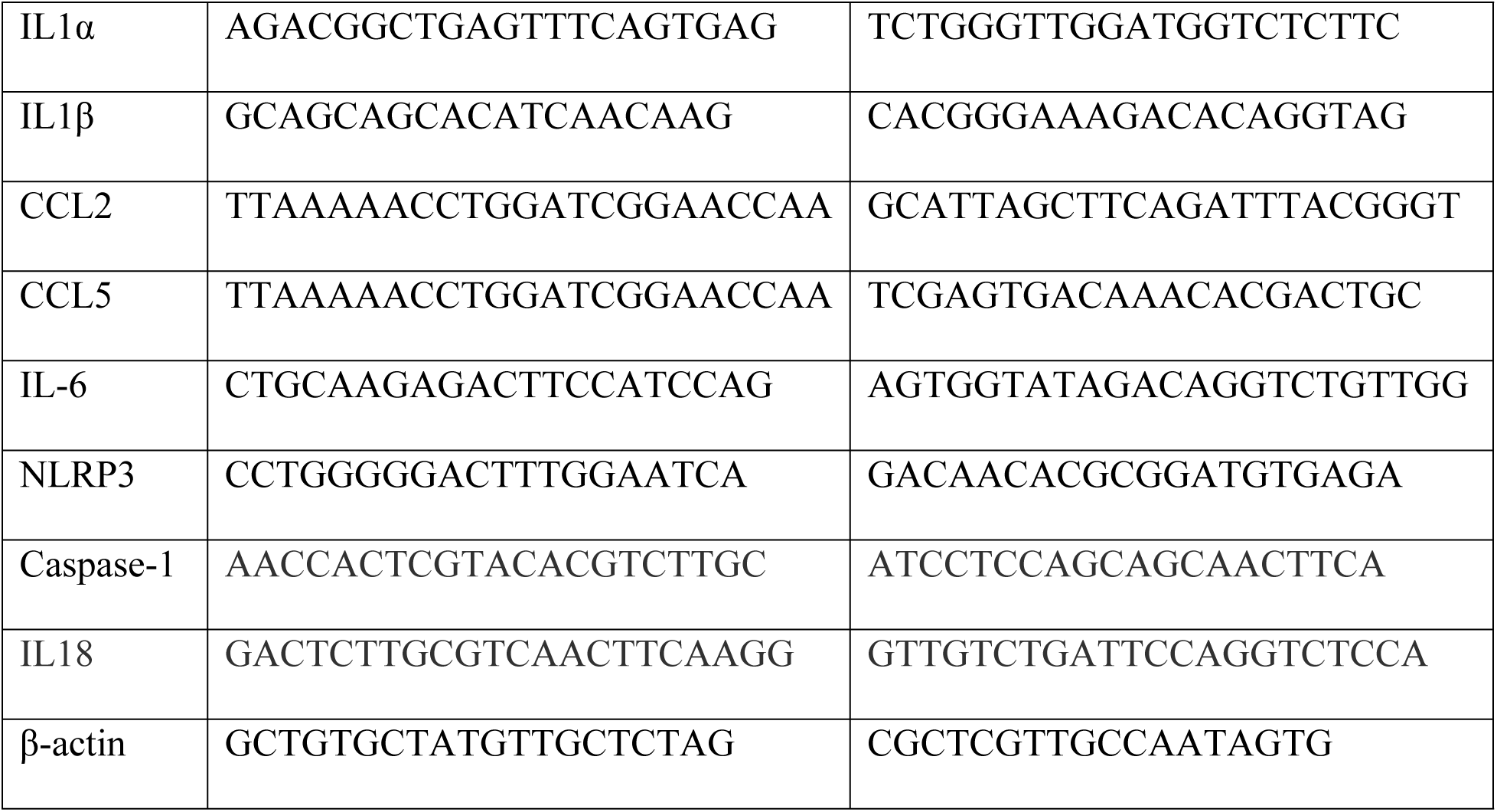
Primer sequences for reverse transcriptase quantitative PCR.

#### Cell Viability Assay

At 7 days post-infection, glial conditioned media (GCM) was removed from mixed glia, centrifuged at 1000 x g for 5 minutes at 4° C to pellet cell debris, and transferred to a fresh tube that was stored at -80° C for viability assays. GCM was thawed in a water bath at 37° C prior to use. N2a neuroblastoma cells were plated on black 96 well Nunc plates (Thermo Fisher Scientific) at 20,000 cells per well. 24 hours after plating, media was removed from N2a cells and replaced with GCM (8-12 replicate wells per treatment group). For each plate, fresh glial media was used as a control for live cells, and glial media containing 0.1% ethanol was used as a control to achieve dead cells. N2as were incubated with GCM or control media for 48 hours. PrestoBlue Cell Viability Reagent (Thermo Fisher Scientific) was allowed to reach room temperature and diluted 1:10 with fresh glial media. Cells were washed gently with PBS and 50 μl PrestoBlue in media was added per well and incubated for 10 minutes at 37° C. Cells were analyzed using the FLUOstar Omega Plate Reader (BMG Labtech). Data was normalized to the live cell control.

#### Scrapie Cell Assay

Primary mixed glial cells were infected with 0.1% RML or normal brain homogenate (NBH), as described above. 7 days later, cells were trypsinized and 50,000 cells were transferred to an ELISpot plate (Millipore). Scrapie cell assay protocol was adapted from Bian et al. 2010 and is described in Hay et al 2022 (24, 74). Primary antibody Sha31 (Cayman Chemical Company) diluted 1:5000 in TBST (Tris-Buffered Saline with Triton-X) was incubated overnight at 4° C. Secondary antibody, AP-α-Mouse IgG (Southern Biotechnology Associates, Birmingham, AL) was diluted 1: 5000 in TBST. Plates were scanned with a ImmunoSpot S6-V analyzer (Cellular Technology Ltd, Shaker Heights, OH), and determined spot numbers using ImmunoSpot5 software (Cellular Technology Ltd, Shaker Heights, OH).

### Brain preparation and inoculation

Brain homogenates were obtained and prepared as described previously (30). Mice were inoculated at 6 weeks of age while under anesthesia with 30μl of 1% Rocky Mountain Laboratories (RML) strain of mouse-adapted scrapie brain homogenate or normal brain homogenate (NBH) for mock-infected animals. Inoculum was prepared in PBS containing 1% PSN.

### Behavioral analyses and clinical scoring

Mice were individually caged at the start of behavioral analysis until euthanasia. Nest building activity was evaluated weekly by placing three fresh napkins in the cage. 48 hours later, the position and structure of the napkins was evaluated, as described previously (30). Nests were evaluated beginning at 13 weeks post-infection (wpi) until euthanasia. Burrowing was analyzed biweekly, as described previously (30), beginning at week 12 and performed every other week, then performed weekly beginning at 16 wpi, for the duration of the study. Clinical signs and weight were evaluated beginning at 13 wpi and continued weekly until mice were euthanized, using previously described scoring methods (30). Clinical signs and weight monitoring were increased to twice weekly once mice began showing clinical scores of 5 or more. Mice were euthanized when they reached a total score of 10 for any combination of signs. Mice inoculated with NBH were used as controls for behavioral analyses and clinical signs. Infected Cx3Cr1Cre-IKKflox mice succumbed to disease around 17 wpi. For direct comparison of tissue, both RML-infected and NBH-treated C57Bl6/J, and NBH-treated Cx3Cr1Cre-IKKflox mice, were euthanized at 17 wpi, despite the non-transgenic wild-type C57Bl6/J and NBH-treated mice not displaying significant clinical signs. Additionally, another cohort of RML-infected C57Bl6/J mice was taken to clinical stages.

### Brain Histology

Animals were deeply anaesthetized with isoflurane before euthanasia by decapitation. Brains were removed and the left hemisphere was frozen at -80° C prior to homogenization. The right hemisphere was fixed in 10% neutral buffered formalin for 72 hours. Tissue was embedded in paraffin and 5 μm sections were cut for staining.

#### Hematoxylin and eosin staining

Hematoxylin and eosin staining of paraffinized brains was performed on the right hemisphere, as described previously (30). Vacuoles in the frontal cortex, hippocampus, thalamus and cerebellum were assessed for quantity and size and scored from 0 (no pathology) to 5 (significant pathology) by three blinded pathologists and averaged.

#### Immunohistochemistry

Immunohistochemical staining of paraffinized brains was performed on the right hemisphere, as described previously (30). Iba1 (Abcam) was used at a 1:400 dilution, GFAP (Dako) was used at a 1:400 dilution, and NeuN (Cell Signaling) was used at a 1:250 dilution. 40x representative images and full brain scans were performed with the Olympus BX53. The Olympus CellSens software (v 1.18) was used to count Iba1+ and GFAP+ cells, and NeuN+ cells in the CA1 region of the hippocampus were counted manually. For NeuN counts, outlier identification and removal were performed using a ROUT outlier test (Q=10%).

#### Immunofluorescence and glial skeletonization

Immunofluorescence and glial skeletonization of paraffinized brains was performed on the right hemisphere, as described previously (30). The following primary antibodies were used: Iba1 (Abcam) at a 1:50 dilution, GFAP (Dako) at a 1:250 dilution, S100β (Abcam) at a 1:750 dilution with C3 (Abcam) at a 1:250 dilution. For each animal, four regions between the dentate gyrus and CA1-CA3 region of the hippocampus were imaged at 40x with an Olympus BX63 fluorescence microscope with a motorized stage and Hamamatsu ORCA-flash 4.0 LT CCD camera and an Olympus Xline apochromat 20X (0.8 N.A.) air objective. Exposures for each stain were kept consistent within each channel. Regions of interest (ROI) were selected with Olympus CellSens software to identify S100β^+^ astrocytes. Mean gray intensity of C3 was determined within the ROI of each S100β^+^ cell containing a visible nucleus. Skeletonization of astrocytes and microglia was performed using IMARIS 9.9.1 using previously described settings (30), with starting point and seed point thresholds adjusted manually.

### Immunoblotting

The left hemispheres were homogenized and protein was quantified with a BCA. Brain homogenates were treated with 2% Sarkosyl, then digested with 100 μg/ml proteinase K (PK) for 1 hour at 37°C, and 5.8 μg protein was loaded per sample. For undigested brain homogenates, 27 μg per brain were loaded. Blots were blocked in 5% non-fat milk buffer and antibodies were diluted in TBS-T. Brain PrP was analyzed with antibodies Sha31 (Cayman Chemical Company) at a 1:10,000 dilution and 12B2 (Wageningen University, Netherlands) at a 1:5000 dilution. Total brain IKKβ was analyzed with antibody IKKβ D3OC6 (Cell Signaling) at 1:1000 dilution and β-actin (Novus) was used as a control at 1:20,000 dilution. All primary antibodies were incubated overnight at 4°C. Cell lysates were isolated using the protein lysis buffer (50mM Tris, 150mM NaCl, 2mM EDTA, 1mM MgCl2, 100mM NaF, 10% glycerol, 1% Triton X-100, 1% Na deoxycholate, 0.1% SDS and 125mM sucrose) supplemented with Phos-STOP and protease inhibitors (Roche). A BCA Protein Assay kit (Thermo Fisher Scientific) was used to quantify protein concentration of lysates, and 500 μg protein was digested with 20 μg/ml PK (Roche) for PrP^Sc^ blots for 1 hour at 37° C. Digestion was terminated with 2mM PMSF and lysates were spun at 40,000 x g for 1 hour at 4° C before being loaded on a gel. For Total PrP, 20 μg protein was used per sample. Samples were run on a 4-20% acrylamide SDS page gels (BioRad) and transferred onto PVDF blotting paper (MilliPore). All other blots were blocked and incubated with antibodies in 5% bovine serum albumin (Sigma-Aldrich). For PrP, primary antibody Bar-224 (Cayman Chemical Company) was used at 1:1,000 dilution. Loading control GAPDH was incubated at a 1:10,000 dilution (MilliPore). The protein antibody complex was visualized using SuperSignal West Pico PLUS Chemiluminescent Substrate (Thermo Fisher Scientific) and visualized with the BioRad ChemiDoc MP. Quantification of average band intensity for cell lysates was performed using the “measure” function on ImageJ. For brain homogenates, the absolute density of each sample was measured and local background subtracted with ImageQuant TL 10.2 analysis software, Cytiva Life Sciences.

### Statistical analysis

Unless stated otherwise, outlier identification and removal was performed using a ROUT outlier test (Q=1%). Cleaned data was measured using a T-test for two groups, or a One-way ANOVA with Tukey’s post-hoc analysis for three or more groups. For survival curves, a Log-rank (Mantel-Cox) test was performed. A p-value of 0.05 or less was considered significant for all analyses. All figures present mean average +/- standard error of the mean (SEM). Prism (v 9.1.0) was used for all data analysis and graph generation.

## Acknowledgements

We want to thank Colorado State University’s Laboratory Animal Resources for providing animal care. We want to thank Tenley French for providing us with mice and Stu Tobet for allowing us to use his IMARIS software, Adam Chicco for allowing us to use his plate reader, and Glenn Telling for allowing us to use his plate scanner.

## References

1. Wang C, Fan L, Khawaja RR, Liu B, Zhan L, Kodama L, et al. Microglial NF-kappaB drives tau spreading and toxicity in a mouse model of tauopathy. Nat Commun. 2022;13(1):1969.

2. Frakes AE, Ferraiuolo L, Haidet-Phillips AM, Schmelzer L, Braun L, Miranda CJ, et al. Microglia induce motor neuron death via the classical NF-kappaB pathway in amyotrophic lateral sclerosis. Neuron. 2014;81(5):1009–23.

3. Rocha SM, Kirkley KS, Chatterjee D, Aboellail TA, Smeyne RJ, Tjalkens RB. Microglia-specific knock-out of NF-kappaB/IKK2 increases the accumulation of misfolded alpha-synuclein through the inhibition of p62/sequestosome-1-dependent autophagy in the rotenone model of Parkinson’s disease. Glia. 2023.

4. Carroll JA, Striebel JF, Race B, Phillips K, Chesebro B. Prion infection of mouse brain reveals multiple new upregulated genes involved in neuroinflammation or signal transduction. J Virol. 2015;89(4):2388–404.

5. Hartmann K, Sepulveda-Falla D, Rose IVL, Madore C, Muth C, Matschke J, et al. Complement 3(+)-astrocytes are highly abundant in prion diseases, but their abolishment led to an accelerated disease course and early dysregulation of microglia. Acta Neuropathol Commun. 2019;7(1):83.

6. Garcao P, Oliveira CR, Agostinho P. Comparative study of microglia activation induced by amyloid-beta and prion peptides: role in neurodegeneration. J Neurosci Res. 2006;84(1):182–93.

7. Kushwaha R, Sinha A, Makarava N, Molesworth K, Baskakov IV. Non-cell autonomous astrocyte-mediated neuronal toxicity in prion diseases. Acta Neuropathol Commun. 2021;9(1):22.

8. Carroll JA, Striebel JF, Rangel A, Woods T, Phillips K, Peterson KE, et al. Prion Strain Differences in Accumulation of PrPSc on Neurons and Glia Are Associated with Similar Expression Profiles of Neuroinflammatory Genes: Comparison of Three Prion Strains. PLoS Pathog. 2016;12(4):e1005551.

9. Kim JI, Ju WK, Choi JH, Choi E, Carp RI, Wisniewski HM, et al. Expression of cytokine genes and increased nuclear factor-kappa B activity in the brains of scrapie-infected mice. Brain Res Mol Brain Res. 1999;73(1-2):17–27.

10. Kim TK, Lee I, Cho JH, Canine B, Keller A, Price ND, et al. Core transcriptional regulatory circuits in prion diseases. Mol Brain. 2020;13(1):10.

11. Bourgognon JM, Spiers JG, Scheiblich H, Antonov A, Bradley SJ, Tobin AB, et al. Alterations in neuronal metabolism contribute to the pathogenesis of prion disease. Cell Death Differ. 2018;25(8):1408–25.

12. Bourteele S, Oesterle K, Weinzierl AO, Paxian S, Riemann M, Schmid RM, et al. Alteration of NF-kappaB activity leads to mitochondrial apoptosis after infection with pathological prion protein. Cell Microbiol. 2007;9(9):2202–17.

13. Julius C, Heikenwalder M, Schwarz P, Marcel A, Karin M, Prinz M, et al. Prion propagation in mice lacking central nervous system NF-kappaB signalling. J Gen Virol. 2008;89(Pt 6):1545–50.

14. Liddelow SA, Guttenplan KA, Clarke LE, Bennett FC, Bohlen CJ, Schirmer L, et al. Neurotoxic reactive astrocytes are induced by activated microglia. Nature. 2017;541(7638):481-7.

15. Hong S, Beja-Glasser VF, Nfonoyim BM, Frouin A, Li S, Ramakrishnan S, et al. Complement and microglia mediate early synapse loss in Alzheimer mouse models. Science. 2016;352(6286):712-6.

16. Guttenplan KA, Weigel MK, Prakash P, Wijewardhane PR, Hasel P, Rufen-Blanchette U, et al. Neurotoxic reactive astrocytes induce cell death via saturated lipids. Nature. 2021;599(7883):102-7.

17. Chen SH, Oyarzabal EA, Sung YF, Chu CH, Wang Q, Chen SL, et al. Microglial regulation of immunological and neuroprotective functions of astroglia. Glia. 2015;63(1):118–31.

18. Srivastava S, Katorcha E, Makarava N, Barrett JP, Loane DJ, Baskakov IV. Inflammatory response of microglia to prions is controlled by sialylation of PrP(Sc). Sci Rep. 2018;8(1):11326.

19. Wardyn JD, Ponsford AH, Sanderson CM. Dissecting molecular cross-talk between Nrf2 and NF-kappaB response pathways. Biochem Soc Trans. 2015;43(4):621–6.

20. Liu T, Zhang L, Joo D, Sun SC. NF-kappaB signaling in inflammation. Signal Transduct Target Ther. 2017;2:17023-.

21. Nuvolone M, Sorce S, Schwarz P, Aguzzi A. Prion pathogenesis in the absence of NLRP3/ASC inflammasomes. PLoS One. 2015;10(2):e0117208.

22. Criollo A, Senovilla L, Authier H, Maiuri MC, Morselli E, Vitale I, et al. The IKK complex contributes to the induction of autophagy. EMBO J. 2010;29(3):619–31.

23. Li ZW, Omori SA, Labuda T, Karin M, Rickert RC. IKK beta is required for peripheral B cell survival and proliferation. J Immunol. 2003;170(9):4630–7.

24. Hay AJD, Murphy TJ, Popichak KA, Zabel MD, Moreno JA. Adipose-derived mesenchymal stromal cells decrease prion-induced glial inflammation in vitro. Sci Rep. 2022;12(1):22567.

25. Easley-Neal C, Foreman O, Sharma N, Zarrin AA, Weimer RM. CSF1R Ligands IL-34 and CSF1 Are Differentially Required for Microglia Development and Maintenance in White and Gray Matter Brain Regions. Front Immunol. 2019;10:2199.

26. Groveman BR, Schwarz B, Bohrnsen E, Foliaki ST, Carroll JA, Wood AR, et al. A PrP EGFR signaling axis controls neural stem cell senescence through modulating cellular energy pathways. J Biol Chem. 2023;299(11):105319.

27. Carroll JA, Chesebro B. Neuroinflammation, Microglia, and Cell-Association during Prion Disease. Viruses. 2019;11(1).

28. Majer A, Medina SJ, Niu Y, Abrenica B, Manguiat KJ, Frost KL, et al. Early mechanisms of pathobiology are revealed by transcriptional temporal dynamics in hippocampal CA1 neurons of prion infected mice. PLoS Pathog. 2012;8(11):e1003002.

29. Moreno JA, Radford H, Peretti D, Steinert JR, Verity N, Martin MG, et al. Sustained translational repression by eIF2alpha-P mediates prion neurodegeneration. Nature. 2012;485(7399):507-11.

30. Hay AJD, Latham AS, Mumford G, Hines AD, Risen S, Gordon E, et al. Intranasally delivered mesenchymal stromal cells decrease glial inflammation early in prion disease. Frontiers in Neuroscience. 2023;17.

31. Hay A, Popichak K, Moreno J, Zabel M. The Role of Glial Cells in Neurobiology and Prion Neuropathology. Cells. 2024;13(10).

32. Sandberg MK, Al-Doujaily H, Sharps B, De Oliveira MW, Schmidt C, Richard-Londt A, et al. Prion neuropathology follows the accumulation of alternate prion protein isoforms after infective titre has peaked. Nat Commun. 2014;5:4347.

33. Wang Y, Hartmann K, Thies E, Mohammadi B, Altmeppen H, Sepulveda-Falla D, et al. Loss of Homeostatic Microglia Signature in Prion Diseases. Cells. 2022;11(19).

34. Gomez-Nicola D, Fransen NL, Suzzi S, Perry VH. Regulation of microglial proliferation during chronic neurodegeneration. J Neurosci. 2013;33(6):2481–93.

35. Liu LR, Liu JC, Bao JS, Bai QQ, Wang GQ. Interaction of Microglia and Astrocytes in the Neurovascular Unit. Front Immunol. 2020;11:1024.

36. Scheckel C, Imeri M, Schwarz P, Aguzzi A. Ribosomal profiling during prion disease uncovers progressive translational derangement in glia but not in neurons. Elife. 2020;9.

37. Aguzzi A, Sigurdson C, Heikenwaelder M. Molecular mechanisms of prion pathogenesis. Annu Rev Pathol. 2008;3:11–40.

38. Lakkaraju AKK, Frontzek K, Lemes E, Herrmann U, Losa M, Marpakwar R, et al. Loss of PIKfyve drives the spongiform degeneration in prion diseases. EMBO Mol Med. 2021;13(9):e14714.

39. Mallucci G, Dickinson A, Linehan J, Klohn PC, Brandner S, Collinge J. Depleting neuronal PrP in prion infection prevents disease and reverses spongiosis. Science. 2003;302(5646):871-4.

40. Lakkaraju AKK, Sorce S, Senatore A, Nuvolone M, Guo J, Schwarz P, et al. Glial activation in prion diseases is selectively triggered by neuronal PrP(Sc). Brain Pathol. 2022;32(5):e13056.

41. Bueler H, Aguzzi A, Sailer A, Greiner RA, Autenried P, Aguet M, et al. Mice devoid of PrP are resistant to scrapie. Cell. 1993;73(7):1339–47.

42. Sorce S, Nuvolone M, Russo G, Chincisan A, Heinzer D, Avar M, et al. Genome-wide transcriptomics identifies an early preclinical signature of prion infection. PLoS Pathog. 2020;16(6):e1008653.

43. Carroll JA, Race B, Williams K, Striebel J, Chesebro B. RNA-seq and network analysis reveal unique glial gene expression signatures during prion infection. Mol Brain. 2020;13(1):71.

44. Moser M, Colello RJ, Pott U, Oesch B. Developmental expression of the prion protein gene in glial cells. Neuron. 1995;14(3):509–17.

45. Victoria GS, Arkhipenko A, Zhu S, Syan S, Zurzolo C. Astrocyte-to-neuron intercellular prion transfer is mediated by cell-cell contact. Sci Rep. 2016;6:20762.

46. Shi F, Yang L, Kouadir M, Yang Y, Wang J, Zhou X, et al. The NALP3 inflammasome is involved in neurotoxic prion peptide-induced microglial activation. J Neuroinflammation. 2012;9:73.

47. Carroll JA, Race B, Williams K, Striebel J, Chesebro B. Microglia Are Critical in Host Defense against Prion Disease. J Virol. 2018;92(15).

48. Race B, Williams K, Baune C, Striebel JF, Long D, Thomas T, et al. Microglia have limited influence on early prion pathogenesis, clearance, or replication. PLoS One. 2022;17(10):e0276850.

49. De Lucia C, Rinchon A, Olmos-Alonso A, Riecken K, Fehse B, Boche D, et al. Microglia regulate hippocampal neurogenesis during chronic neurodegeneration. Brain Behav Immun. 2016;55:179–90.

50. Zhu C, Herrmann US, Falsig J, Abakumova I, Nuvolone M, Schwarz P, et al. A neuroprotective role for microglia in prion diseases. J Exp Med. 2016;213(6):1047–59.

51. Bradford BM, McGuire LI, Hume DA, Pridans C, Mabbott NA. Microglia deficiency accelerates prion disease but does not enhance prion accumulation in the brain. Glia. 2022;70(11):2169–87.

52. Shan Z, Hirai Y, Nakayama M, Hayashi R, Yamasaki T, Hasebe R, et al. Therapeutic effect of autologous compact bone-derived mesenchymal stem cell transplantation on prion disease. J Gen Virol. 2017;98(10):2615–27.

53. Song CH, Honmou O, Ohsawa N, Nakamura K, Hamada H, Furuoka H, et al. Effect of transplantation of bone marrow-derived mesenchymal stem cells on mice infected with prions. J Virol. 2009;83(11):5918–27.

54. de Melo A, Lima JLD, Malta MCS, Marroquim NF, Moreira AR, de Almeida Ladeia I, et al. The role of microglia in prion diseases and possible therapeutic targets: a literature review. Prion. 2021;15(1):191–206.

55. Shi F, Yang L, Wang J, Kouadir M, Yang Y, Fu Y, et al. Inhibition of phagocytosis reduced the classical activation of BV2 microglia induced by amyloidogenic fragments of beta-amyloid and prion proteins. Acta Biochim Biophys Sin (Shanghai). 2013;45(11):973–8.

56. Chan WY, Kohsaka S, Rezaie P. The origin and cell lineage of microglia: new concepts. Brain Res Rev. 2007;53(2):344–54.

57. Saura J. Microglial cells in astroglial cultures: a cautionary note. J Neuroinflammation. 2007;4:26.

58. Carroll JA, Race B, Williams K, Striebel JF, Chesebro B. Innate immune responses after stimulation with Toll-like receptor agonists in ex vivo microglial cultures and an in vivo model using mice with reduced microglia. J Neuroinflammation. 2021;18(1):194.

59. He M, Dong H, Huang Y, Lu S, Zhang S, Qian Y, et al. Astrocyte-Derived CCL2 is Associated with M1 Activation and Recruitment of Cultured Microglial Cells. Cell Physiol Biochem. 2016;38(3):859–70.

60. Liddelow SA, Barres BA. Reactive Astrocytes: Production, Function, and Therapeutic Potential. Immunity. 2017;46(6):957–67.

61. Jack CS, Arbour N, Manusow J, Montgrain V, Blain M, McCrea E, et al. TLR signaling tailors innate immune responses in human microglia and astrocytes. J Immunol. 2005;175(7):4320–30.

62. Holm TH, Draeby D, Owens T. Microglia are required for astroglial Toll-like receptor 4 response and for optimal TLR2 and TLR3 response. Glia. 2012;60(4):630–8.

63. Carroll JA, Race B, Williams K, Chesebro B. Toll-like receptor 2 confers partial neuroprotection during prion disease. PLoS One. 2018;13(12):e0208559.

64. Spinner DS, Cho IS, Park SY, Kim JI, Meeker HC, Ye X, et al. Accelerated prion disease pathogenesis in Toll-like receptor 4 signaling-mutant mice. J Virol. 2008;82(21):10701–8.

65. Chhor V, Le Charpentier T, Lebon S, Ore MV, Celador IL, Josserand J, et al. Characterization of phenotype markers and neuronotoxic potential of polarised primary microglia in vitro. Brain Behav Immun. 2013;32:70–85.

66. Shi F, Yang Y, Kouadir M, Fu Y, Yang L, Zhou X, et al. Inhibition of phagocytosis and lysosomal acidification suppresses neurotoxic prion peptide-induced NALP3 inflammasome activation in BV2 microglia. J Neuroimmunol. 2013;260(1-2):121–5.

67. Sigurdson CJ, Bartz JC, Glatzel M. Cellular and Molecular Mechanisms of Prion Disease. Annu Rev Pathol. 2019;14:497–516.

68. Cortes CJ, Qin K, Cook J, Solanki A, Mastrianni JA. Rapamycin delays disease onset and prevents PrP plaque deposition in a mouse model of Gerstmann-Straussler-Scheinker disease. J Neurosci. 2012;32(36):12396–405.

69. Xu Y, Zhang J, Tian C, Ren K, Yan YE, Wang K, et al. Overexpression of p62/SQSTM1 promotes the degradations of abnormally accumulated PrP mutants in cytoplasm and relieves the associated cytotoxicities via autophagy-lysosome-dependent way. Med Microbiol Immunol. 2014;203(2):73–84.

70. Shim SY, Karri S, Law S, Schatzl HM, Gilch S. Prion infection impairs lysosomal degradation capacity by interfering with rab7 membrane attachment in neuronal cells. Sci Rep. 2016;6:21658.

71. Ma S, Attarwala IY, Xie XQ. SQSTM1/p62: A Potential Target for Neurodegenerative Disease. ACS Chem Neurosci. 2019;10(5):2094–114.

72. Chen Y, Li Q, Li Q, Xing S, Liu Y, Liu Y, et al. p62/SQSTM1, a Central but Unexploited Target: Advances in Its Physiological/Pathogenic Functions and Small Molecular Modulators. J Med Chem. 2020;63(18):10135–57.

73. Livak KJ, Schmittgen TD. Analysis of relative gene expression data using real-time quantitative PCR and the 2(-Delta Delta C(T)) Method. Methods. 2001;25(4):402–8.

74. Bian J, Napier D, Khaychuck V, Angers R, Graham C, Telling G. Cell-based quantification of chronic wasting disease prions. J Virol. 2010;84(16):8322–6.

